# A functional artificial neural network for noninvasive presurgical evaluation of glioblastoma multiforme prognosis and radiosensitivity profiling

**DOI:** 10.1101/2020.12.15.422749

**Authors:** Eric Zander, Andrew Ardeleanu, Ryan Singleton, Barnabas Bede, Yilin Wu, Shuhua Zheng

**Affiliations:** DigiPen Institute of Technology, Department of Mathematics, Washington, 98052, U.S.A.; Nova Southeastern University, College of Osteopathic Medicine, Florida, 33314, U.S.A.

**Keywords:** cullin2, glioblastoma multiforme, copy number variations, artificial intelligence, machine learning, deep learning

## Abstract

**Background and Purpose:** Genetic profiling for glioblastoma multiforme (GBM) patients with intracranial biopsy carries a significant risk of permanent morbidity. We previously demonstrated that the *CUL2* gene, encoding the scaffold cullin2 protein in the cullin2-RING E3 ligase (CRL2), can predict GBM radiosensitivity and prognosis mainly due to the functional involvement of CRL2 in mediating hypoxia-inducible factor 1 (HIF-1) α and epidermal growth factor receptor (EGFR) degradation. Because *CUL2* expression levels are closely regulated with its copy number variations (CNVs), this study aims to develop an artificial neural network (ANN) that can predict GBM prognosis and help optimize personalized GBM treatment planning.

**Materials and Methods:** Datasets including Ivy-GAP, The Cancer Genome Atlas Glioblastoma Multiforme (TCGA-GBM), the Chinese Glioma Genome Atlas (CGGA) were analyzed. T1 images from corresponding cases were studied using automated segmentation for features of heterogeneity and tumor edge contouring.

**Results:** We developed a 4-layer neural network that can consistently predict GBM prognosis with 80-85% accuracy with 3 inputs including *CUL2* copy number, patient’s age at GBM diagnosis, and surface vs. volume (SvV) ratio.

**Conclusion:** A functional 4-layer neural network was constructed that can predict GBM prognosis and potential radiosensitivity.

## Introduction

Glioblastoma Multiforme (GBM) is an aggressive form of tumor in the central nervous system (CNS) with less than 5% of patients in the United States surviving for five years following initial diagnosis[1]. While concurrent and adjuvant chemoradiotherapy constitutes standardized treatment for GBM after the Stupp *et al*. clinical trial in 2005, finding further methods of refining treatment would prove invaluable for better patient outcomes with less general toxicity[2]. Recent research regarding biomarkers of traits governing treatment response as well as promising machine learning (ML) applications in neuroimaging analysis join to indicate that targeted treatment informed by genetic and imaging data could lead to the paradigm shift necessary for such improvement[3, 4]. The rapidly developing field of ML includes methods allowing for increasingly robust approaches to research designed to enhance targeted treatment. Aspects of deep learning (DL), a subfield of ML concerned with multi-layered artificial neural networks (ANNs), stand to benefit GBM research in particular given that applications in medical image processing allow the systematic extraction of image features desirable for use in predictive modeling [4-6].

Using genetic data from numerous datasets, we recently demonstrated expression levels of *CUL2*, which encodes the cullin2 scaffold protein the cullin2-RING E3 ligase (CRL2) complex, can predict GBM radiosensitivity[7]. Variations of GBM radiosensitivity has been well demonstrated in xenograft models with single-dose irradiation TCD 50s (tumor control dose 50%) values range from 32.5 to 75.2 Gy[8, 9], indicating that the ability to predict a GBM patient’s radiosensitivity could promote effective personalized RT and improved patient outcomes as a result. Notably, *CUL2* copy number variation (CNV) dictates expression levels, suggesting that models could rely on CNVs measured through non-invasive methods rather than expression levels to predict a patient’s radiosensitivity[7, 10]. To improve the accuracy of these predictions without requiring additional invasive techniques, ANN models could also leverage data extracted from MRI to help assess treatment response and patient prognosis[11]. Neuroimaging data contains valuable information for measuring treatment response as well as discerning patient outlook. Distinct quantitative GBM image features such as tumor shape, edge sharpness, and texture varied with the survival probability of patients[12]. Combined with clinical measurements found in existing datasets, features extracted from imaging data could be combined with genetic information such as *CUL2* CNVs to strengthen the foundation for non-invasive, effective, and consistent evaluation of treatment outcomes and radiosensitivity profiling.

To this end, we employed ML and DL methodologies to create an effective model for radiosensitivity profiling using *CUL2* CNVs and imaging data derived from independent public datasets. An ANN working model that integrates image, clinical, and genetic information for non-invasive radiosensitivity profiling is proposed.

## Materials and Methods

### Public Datasets

Datasets including genetic data regarding *CUL2* copy number variations (CNVs) and expression levels, clinical data indicating patient demographic and overall survival (OS), and T1 Magnetic Resonance Imaging (MRI) images enabled this study. Such datasets include The Cancer Genome Atlas Glioblastoma Multiforme (TCGA-GBM), the Ivy Glioblastoma Atlas Project (Ivy-GAP), and the Chinese Glioma Genome Atlas (CGGA)[13-15]. Clinical and genetic data from the TCGA-GBM were acquired through the Xena platform (https://xena.ucsc.edu/)[16]. TCGA-GBM images were made available through The Cancer Imaging Archive (TCIA)[17]. This also includes skull-stripped and co-registered segmentations of TCGA-GBM images made available by Bakas *et al*., as well as DICOM-SEG conversions of these segmented images created by Beers *et al*.[17]. Images from the Ivy-GAP dataset were also acquired through TCIA, with clinical and genetic data made available by Puchalski *et al*[14].

### T1 MRI image segmentation

While images considered in this research included post-gadolinium T1-weighted DICOM images from the original TCGA-GBM dataset, the features used in modeling were derived from the most voluminous DICOM-SEG conversions of images in the segmented image dataset for each patient. Python packages for processing image files in the DICOM, DICOM-SEG, and NIfTI file formats include pydicom, pydicom_seg, and nibabel, respectively.

### Image Analyses

To explore the applicability of image features for the prediction of patient outcomes, a simple ratio comparing the surface area of tumor borders to the total volume of borders was derived from the DICOM-SEG conversions of segmented tumors. The edges of these borders can be detected using the canny edge detector such as the one available from the Skimage, or Scikit-image, Python package. Finding the sum of the pixels forming these edges gives a close approximation of surface area. Tumor border volume can be calculated by summing up all pixels within the tumor borders. Calculating the tumor surface area vs. volume (SvV) ratio between the two is then as simple as dividing the surface area by volume (SvV = Surface Area / Volume). This results in a value unique to each patient that is indicative of tumor border regularity[12].

### Kaplan–Meier (K–M) survival analysis

K–M survival analyses for GBM with differential *CUL2* copy numbers, Karnofsky Performance Score (KPS) rankings, age at GBM diagnosis, surface area, and SvV ratios were conducted using the Lifelines library. The data set is split into two at the mean value of any of respective attributes above. Any value that is above the mean value for that attribute is placed in the upper group, while any value that is below the mean is placed in the lower group. Therefore, the number of patients in either group varies from attribute to attribute. Because of this, the number of patients in any one group is labeled next to the legend for that group on its graph. Log-Rank analyses for p-values smaller than 0.05 were considered statistically significant.

### Artificial neural network (ANN) of GBM genomics and clinical features

Four different ANNs were created for the purpose of predicting the OS time of patients with the goal of testing whether or not measuring patient *CUL2* copy numbers would yield similar results to *CUL2* expression levels. Results are determined by how often a neural network can predict a patient’s survivability based on clinical data. Four different loss functions were used to test on all four of our neural networks, resulting in 16 different results to compare. The loss functions we chose were Binary Crossentropy, Mean Absolute Error, Mean Error Squared, and Categorical Crossentropy. The packages used for creating these neural networks include Keras, Pytorch, and TensorFlow. After extensive testing, a 4-layer model (8-8-8-2), with a binary output, was built.

The first layer consists of 8 nodes and takes number of inputs based on the information we are passing to it. The information being passed to our neural networks is as follows: Baseline (Age, KPS, longest dimension), Expression (*CUL2* expression, Age, KPS, longest dimension), Copy Number Variation (*CUL2* copy numbers, Age, KPS, longest dimension), and Feature Data (*CUL2* copy numbers, Age, SvV). Therefore, each neural network requires a different amount of inputs. The second and third layer are both also 8 nodes and use a relu activation function. The last layer is binary output, with outputs of either 0 or 1.

Regarding the last binary output layer of the neural network, we split each dataset into targets of 0 or 1, with patients who are assigned a 1 survived longer, and those who did not are assigned a 0. The data is split to be approximately 55%-45% in favor of 0 targets, with the exception of the feature dataset which is split approximately 60%-40% in favor of 0 targets. The data must be split like this in order to ensure a valid model. **Figure 1** is a schematic overview of the architecture of the ANN. The data is passed into each of the 8 nodes, and so on until the last output layer, and if the model is correct it reinforces the model, whereas if it is wrong it punishes the model and tweaks the weights of each neuron accordingly.

**Fig. 1.**
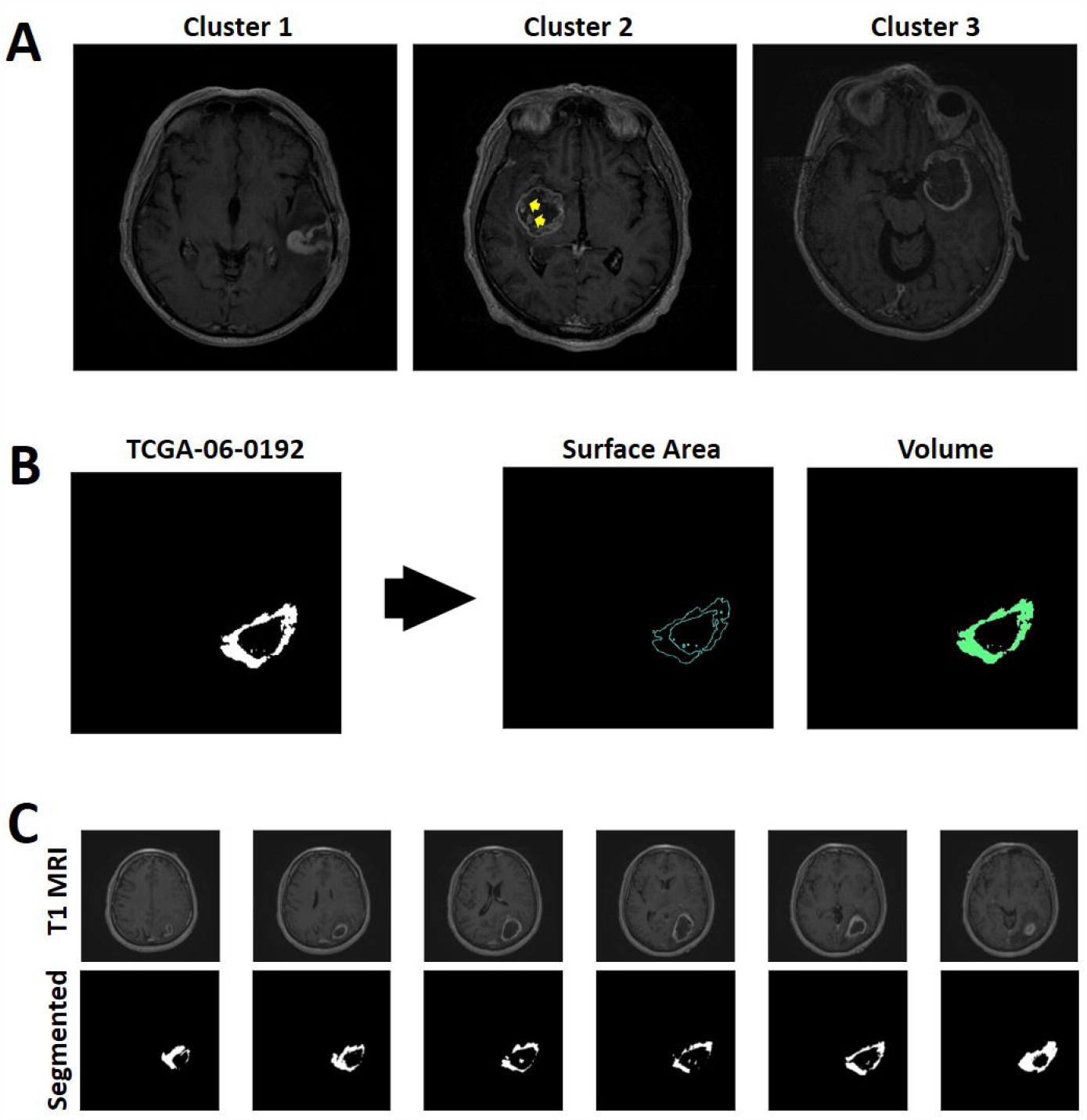
GBM T1 image segmentation. **A)** Example of imaging phenotypes as defined by Itakura et al.[12] with multifocal Cluster 1, spherical Cluster 2, and rim-enhancing Cluster 3 GBM cases. Arrows indicate intralesional multifocal heterogeneity. **B)** Example of binary segmentation masks attainable from DICOM-SEG (DSO) conversions for the TCGA-GBM dataset. The surface area of tumor borders calculated by summing up edges of 2D slices derived with a canny edge detector. The volumes were approximated by summing up all pixels of each slice. **C)** Examples of segmented images with corresponding T1 image from the GBM case TCGA-06-0192.

## Results

### GBM T1 MRI segmentation and tumor surface regularity

Many radiomic features such as first order statistics measuring gray value intensity and skewness, texture features describing the tumor region serve as predictors for GBM overall survival (OS) rates[18]. One method of systematically measuring shape involves finding the surface area vs. volume ratio (SvV) of tumor borders[19]. Using the most voluminous binary segmentation masks attainable from DICOM-SEG (DSO) conversions for the TCGA-GBM segmentation dataset for each patient, the surface area of tumor borders was approximated by summing up edges of 2D slices derived with a canny edge detector (**Figure 1B**). The volumes of segmented tumors masks can then be calculated by summing up all pixels of each slice. Applying this approach on the segmented image dataset creates surface area, volume, and SvV ratio data for 102 patients, 98 of these patients have recorded *CUL2* CNV averages in the corresponding genetic dataset.

Itakura *et al*. established three clusters of GBM phenotypes associated with distinct OS categories[12]. These clusters are distinguished by concavity and regional intensity with Cluster 1 described as pre-multi-focal tumors that have irregular tumor shapes combined with concavities along their border and associated with poor OS rates; Cluster 2 as spherical with better survival outcomes; Cluster 3 tumors resemble spherical tumors with rim-enhancing and cystic hypointense centers (**Figure 1A**)[12]. However, measuring the outer surface area of GBM was unable of reflect the intralesional heterogeneous features including multifocal hemorrhage and masses, cystic and necrotic components (**Figure 1A**, arrows). By using binary segmented masks, we were able to quantify the GBM intralesional heterogeneity which may help differentiate the Cluster 2 spherical tumors from Cluster 3 rim-enhancing tumors (**Figure 1A, B, C**). Notably, the orientation of these segmentation masks does not affect these measurements (**Figure 1C**). The resulting ratios prove unique to patients and indicate surface regularity.

### Segmented image features

To further investigate the clinical relevance of proposed segmentation methods, we studied the prognostic value of quantifiable image features in GBM. These features include the total surface area, tumor volume, SvV ratio, and longest tumor dimension (**Figure 2**). In Kaplan Meier analyses, we found GBM total surface area based on the segmented images can predict GBM prognosis with patient with lower surface area (n=58) have significantly better prognosis than those with higher surface area (n=39) (*p*=0.0016) (**Figure 2A**). Interestingly, we did not observe significant difference on survival when patients were grouped based on tumor volume, longest dimension, or SvV ratio (**Figure 2B, C, D**). Given that each parameter in the MRI images is unique for individual GBM patient, we tested and tried these image features to find best candidates that can be incorporated for best performance in our neural network.

**Fig. 2.**
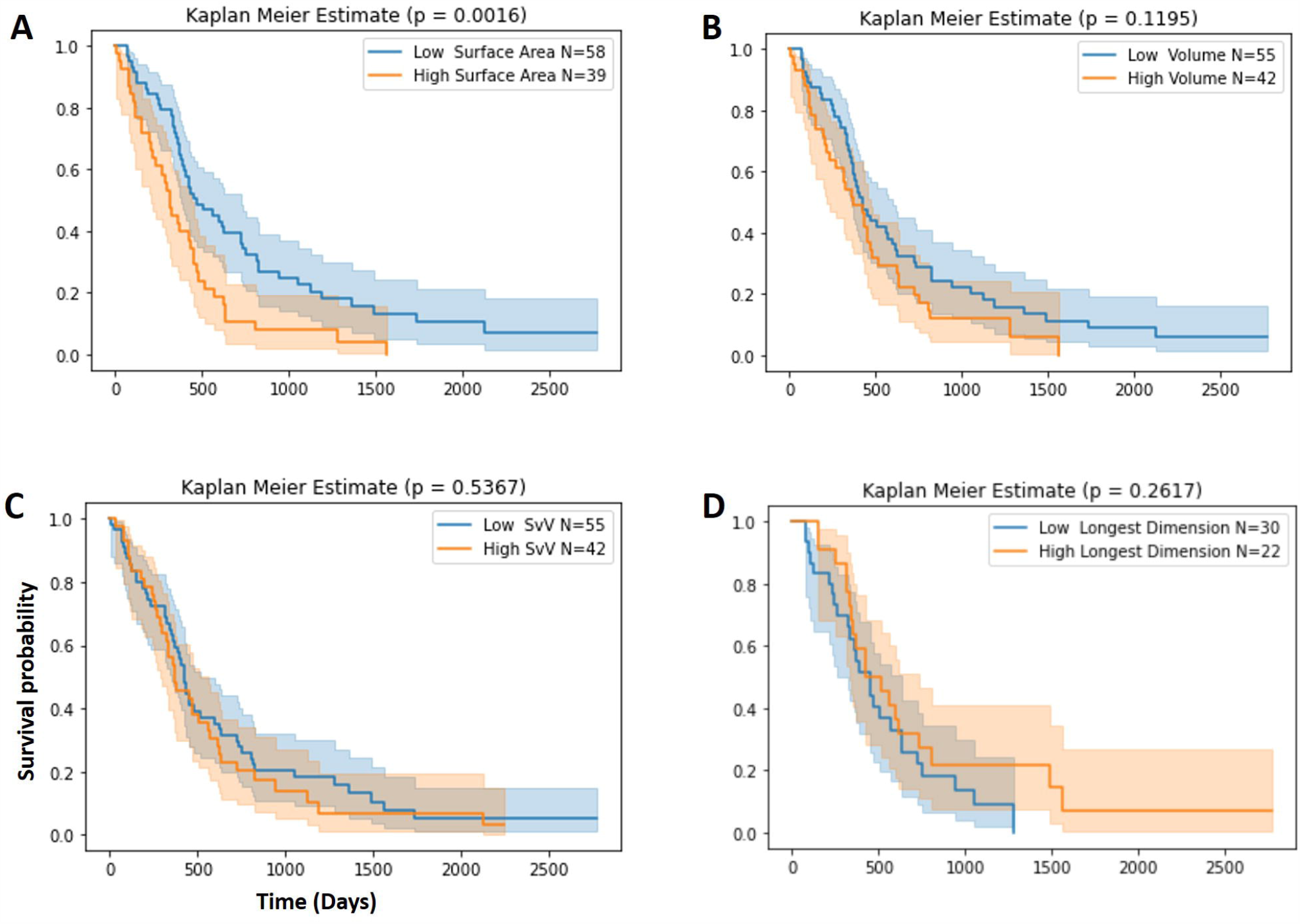
Prognostic value of image features. **A, B, C)** Overall survival (OS) analysis of GBM cases based on the quantifiable features including Surface area, Volume, and Surface area vs. Volume (SvV) of segmented images. Mean values were used as cutoffs for each survival analysis panel. The number of patients in each group was indicated as ‘N=’. **D)** OS analysis of GBM cases from TCGA-GBM dataset based on the longest dimension of GBM in T1 images. Mean value was used as cutoff. The number of patients in each group was indicated as ‘N=’.

### *CUL2* CNVs and GBM surface heterogeneity

We previously demonstrated elevated *CUL2* expression correlates with lower cellular protein levels of hypoxia-inducible factor 1 α (HIF-1 α) and epidermal growth factor receptor (EGFR) due to the polyubiquitination activity of cullin2-RING E3 ligase (CRL2) against these substrate proteins[7, 20, 21]. The importance of HIF-1 α and EGFR in GBM intralesional and surface heterogeneity may correlate *CUL2* expression levels/copy numbers with SvV ratios[22, 23]. We found GBM patients with higher *CUL2* copy numbers are more likely to have spherical or rim-enhancing tumors (Cluster 2, 3, respectively) with corresponding larger surface area as demonstrated in images showing segmentation masks (**Figure 3A**). A ‘left shift’ distribution pattern for Volume and Surface Area with low *CUL2* copy numbers were observed as compared with cases of high *CUL2* copy numbers (**Figure 3B**). However, no significant difference on average Volume, Surface Area and SvV ratio were identified in cases with low and high *CUL2* copy numbers (**Supplementary Figure 1**).

**Fig. 3.**
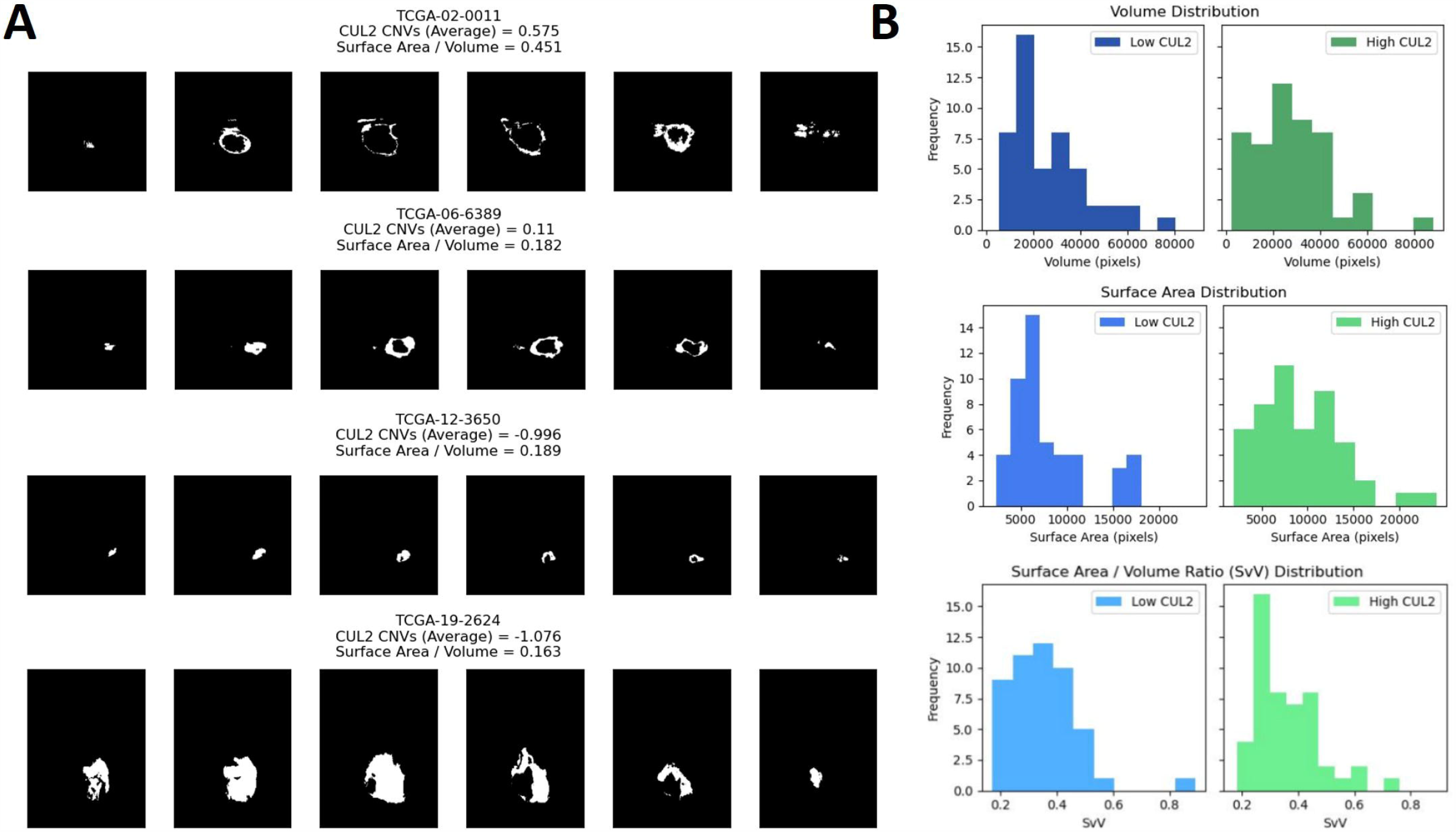
Radiogenomic study of *CUL2* copy numbers and segmented T1 images. **A)** Representative of segmented images of GBM cases with differential *CUL2* copy numbers. **B)** Distribution of Volume, Surface area and Surface area vs. Volume (SvV) in GBM cases with differential *CUL2* copy numbers.

### Prognostic value of *CUL2* CNVs and clinical features

Consistent with what we have reported, GBM patients with higher *CUL2* copy numbers (n=41) often have better OS rates that those with lower *CUL2* copy numbers (n=44) (**Figure 4B**). Given that *CUL2* CNVs and GBM surface area and volume are features that can be obtain via non-invasive methods, we went further to investigate the prognostic value of clinical parameters that are readily available at initial disease diagnosis. Age refers to a patient’s age at initial diagnosis. Patients were split into two groups at their mean age. Patients over the mean age fall into the “High Age” group (n=46), and those that fall under the mean age fall into the “Low Age” group (n=39) (*p*=0.0062) (**Figure 4A**). And as expected those who rank higher (n=68) on the Karnofsky Performance Scale (KPS) tend to survive longer that those with lower ranking (n=17) (**Figure 4C**). We then chose these clinical attributes and *CUL2* CNVs as inputs in our neural network.

**Fig. 4.**
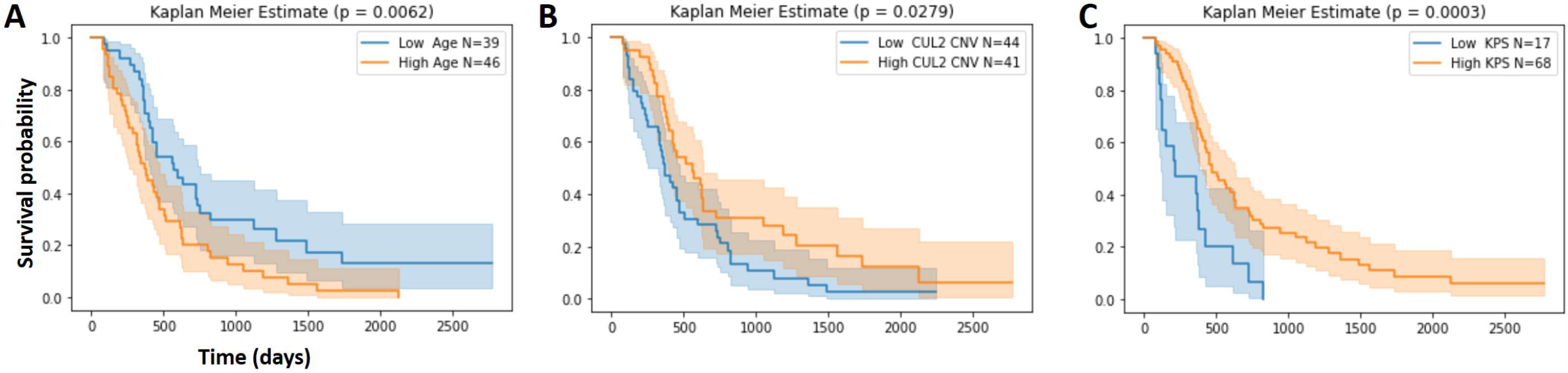
Impact of *CUL2* CNVs and clinical parameters in GBM survival. **A, B, C)** Overall survival (OS) analyses of GBM cases with differential *CUL2* copy numbers, age at GBM diagnosis, and Karnofsky Performance Score (KPS) rankings. Mean values in each panel were used as cutoffs in grouping patients.

### Constructed artificial neural networks (ANNs) predict GBM survival

The parameters that we chose to build the neural networks should stay consistent throughout our trials. A loss function is required in many neural networks and can lead to performance increases/decreases depending on the type of problem that is being tackled[24]. Because of this we employed 4 different loss functions to test on all 4 of our neural networks, resulting in 16 different results to compare. The loss functions we chose were Binary Crossentropy, Mean Absolute Error, Mean Error Squared, and Categorical Crossentropy (**Figure 5A**)[24]. The structure of the neural network we chose is a 4-layer model (8-8-8-2), with a binary output. The first layer consists of 8 nodes with sets of inputs as follows: Baseline dataset (Age, KPS, longest dimension), Expression dataset (*CUL2* expression, Age, KPS, longest dimension), Copy Number Variation dataset (*CUL2* copy numbers, Age, KPS, longest dimension), and Feature Data dataset (*CUL2* copy numbers, Age, SvV) (**Figure 5A**). The second and third layer are both also 8 nodes with a relu activation function. The last layer is binary output, with outputs of either ‘0’ or ‘1’ with patients who are assigned a ‘1’ survived longer, and those who did not are assigned a ‘0’ (**Figure 5C**).

**Fig. 5.**
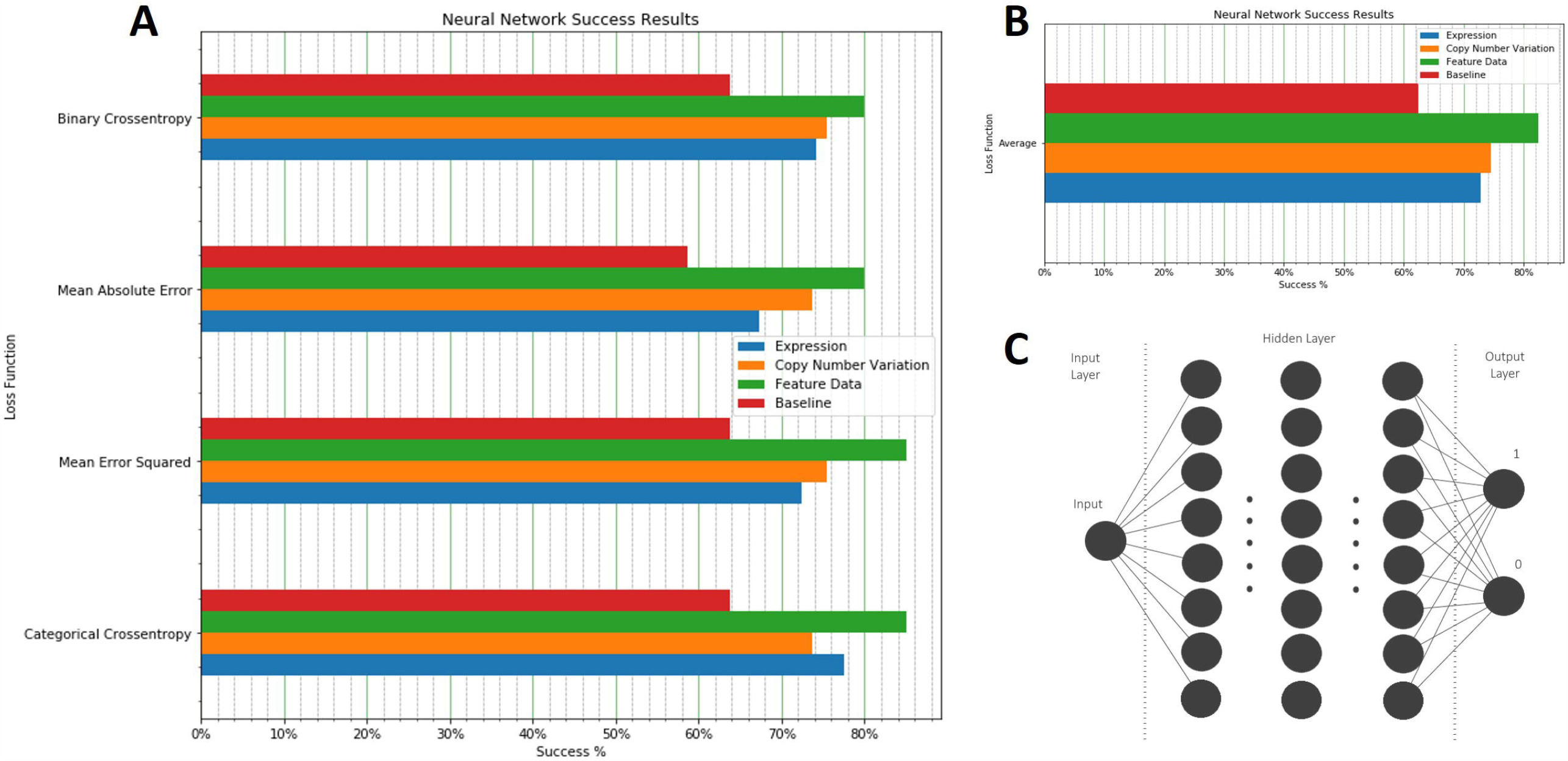
Parameterizing artificial neural networks (ANNs) based on *CUL2* CNVs, image features and clinical data. **A)** 4 loss functions analyses including Binary Crossentropy, Mean Absolute Error, Mean Error Squared, and Categorical Crossentropy were used in our ANNs. The packages used for creating these ANNs include Keras, Pytorch, and TensorFlow. The information being passed to our neural networks is as follows: Baseline (Age, KPS, longest dimension), Expression (*CUL2* expression, Age, KPS, longest dimension), Copy Number Variation (*CUL2* copy numbers, Age, KPS, longest dimension), and Feature Data (*CUL2* copy numbers, Age, SvV). Each neural network run 1000 times, and the prediction results from the best performing model of these 1000 runs were presented. A total of 16 neural networks were trained with a total of 16,000 trials. **B)** Average performance of 4 neural networks for each set of inputs. **C)** A schematic overview of our 4-layer neural networks.

As expected, since the Baseline neural network was given the least amount of data, it makes sense that it was the poorest performing model in all four trials (**Figure 5A**). We then studied the performance between the Copy Number Variation neural network vs. the Expression neural network. The CNV neural network outperformed or similar to the Expression neural network in all four trials (**Figure 5A**). Therefore, *CUL2* expression levels or *CUL2* copy numbers will yield similar results, with around 73-74% accuracy as displayed on the average of the 4 loss functions combined in a single visual (**Figure 5B**).

Lastly, to identify the set of inputs that establish best performing neural network, we have a set number of trials for each of our 16 neural networks. Each neural network run 1000 times with a total of 16,000 trials for 16 neural networks. The best performing model for each set of dataset input was saved for distribution and to view the model’s structure and weights. We found a consistent 80-85% accuracy on all loss functions for Feature Data dataset that has inputs of *CUL2* copy numbers, Age, and SvV (**Figure 5A**).

## Discussion

We previously demonstrated that *CUL2* gene expression levels and copy number variations (CNVs) can predict GBM radiosensitivity and overall survival (OS). This study further constructed neural networks via deep learning (DL) of basic clinical information, T1 MRI-based imaging features and *CUL2* copy numbers. In our best performing neural network models, we consistently demonstrated 80-85% accuracy in predicting GBM prognosis with inputs of *CUL2* copy number, patient’s age at GBM diagnosis, and surface vs. volume (SvV) ratio in segmented images. All these inputs are objective quantifiable parameters that can be obtained without any intracranial biopsy which carries significant risk of causing serious permanent morbidity (5%)[25]. Therefore, our model provides a unique tool for non-invasive pre-surgical evaluation of GBM patients regarding prognosis and potentially radiosensitivity.

We found the usage of tumor border surface area and volume led to improved accuracy for each neural network model. This not only suggests that the uniqueness of these measurements has enough of a degree of influence on OS rates to clearly impact the results, but also demonstrates the general value of quantifiable features derived from image processing in predicting treatment outcomes. In addition to including other measurements of shape indicating surface regularity, future models could also potentially integrate first order statistics regarding gray level intensity, textural features, and a myriad of radiomic features with demonstrable impact on GBM survival. Artificial intelligence-based automated image segmentation will play a critical role in this aspect.

Processing medical images such as T1 images has its challenges, but several approaches exist to extract hundreds of potentially useful features of each category. Image processing solutions that employ convolutional neural network architectures such as 3D U-Net to automate tumor segmentation allow the systematic tumor segmentation, though atlas. Ultimately, additional image features derived using deep learning (DL) solutions stand to improve the already promising model by introducing more non-invasive data alongside *CUL2* CNVs, further indicating the value of combining such measurements for predicting treatment response.

Although we built a promising model for non-invasive pre-surgical evaluation of GBM patients, this study has several limitations. First, this study needs to incorporate further imaging data and features from independent datasets into our neural network which will help our existing neural networks more accurately predict survivability. Second, the radiosensitivity of GBM patients is difficult to reflect with survival data that are derived from retrospective studies. Clinical trials are warranted to fully validate whether the neural network we built can guide RT treatment planning. Furthermore, relying on clinical data can be a risk, especially when certain clinical parameters are subjective. To solve this issue, we need to create new neural networks that are only fed imaging data and patients’ age, without including additional clinical data. Current models we built have binary outputs (‘1’ and ‘0’). Instead of the neural network just trying to predict between two possible outcomes, we could have the neural network try and be more accurate and predict possibly three or four possible outcomes.

