## Supplementary Figure 1 for "A functional artificial neural network for noninvasive presurgical evaluation of glioblastoma multiforme prognosis and radiosensitivity profiling"

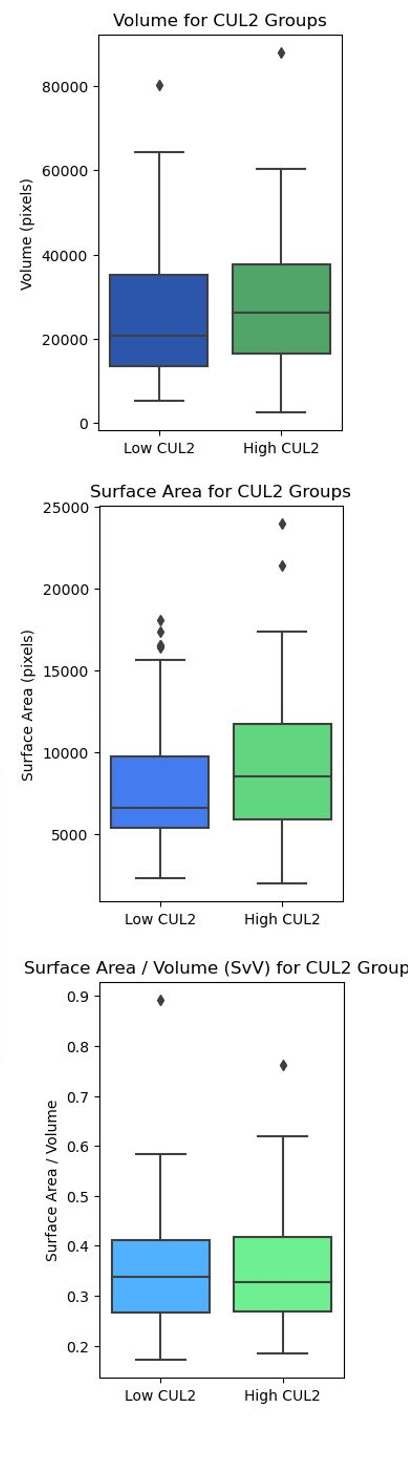

**Suppl. Fig. 1** Distribution of GBM tumor Volume, Surface area, and Surface vs. Volume ratio in cohorts with differential *CUL2* copy numbers.
